# Fragile nucleosomes are essential for RNA Polymerase II to transcribe in eukaryotes

**DOI:** 10.1101/2025.10.08.681188

**Authors:** Lingbo Li, Samuel Hunter, Sonia Leach, Yonghua Zhuang, Haolin Liu, Junfeng Gao, Qianqian Zhang, Timothy J. Stasevich, Hiroshi Kimura, Robin Dowell, Gongyi Zhang

## Abstract

Nucleosomes are barriers to RNA Polymerase II (Pol II) transcribing along gene bodies in eukaryotes. We found that a fragile “tailless nucleosome” could be generated to resolve the Pol II pausing caused by the +1 nucleosome in metazoans. We hypothesized that “pan-acetylated nucleosomes” along gene bodies are fragile and are generated to resolve the resistance of nucleosomes to the elongation of Pol II. To confirm this hypothesis, we systematically analyzed pan-acetylation and pan-phosphorylation of the entire human genome in HEK293T cells. Herein, we show that both pan-acetylation and pan-phosphorylation of histones in nucleosomes along gene bodies are coupled with the activities of Pol II. Our results show that there are four major types of fragile nucleosomes required for Pol II to transcribe along gene bodies: nucleosomes without tails, nucleosomes with dominant acetylation, nucleosomes with dominant phosphorylation, and nucleosomes with both acetylation and phosphorylation.

## Introduction

RNA polymerases (RNAPs) are functionally and structurally conserved from bacteria to human beings (Roeder and Rutter 1969, Zhang, Campbell et al. 1999, Cramer, Bushnell et al. 2000, Landick 2006). In bacteria, transcription can be accomplished by transcription factors, naked DNA, sigma factors, and RNAPs (Burgess and Anthony 2001). In eukaryotes, in reconstituted minimal transcription systems *in vitro,* Pol II can act similarly, transcribing naked DNA with the help of only a few general transcription factors (Matsui, Segall et al. 1980, Samuels, Fire et al. 1982, Usuda, Kubota et al. 1991, Goodrich and Tjian 1994). However, nucleosomes become the barriers to Pol II transcription in eukaryotes (Kornberg 1974, Kornberg and Thomas 1974, Olins and Olins 1974). From in vitro assays, even a single nucleosome could block the transcription RNAPs from phages or Pol II from eukaryotes (Izban and Luse 1991, Kireeva, Walter et al. 2002, Kireeva, Hancock et al. 2005). In higher eukaryotes, only + 1 nucleosome could cause promoter-proximal Pol II pausing for a large group of genes in vivo (Mavrich, Jiang et al. 2008, Schones, Cui et al. 2008, Weber, Ramachandran et al. 2014). RNAP could transcribe smoothly on tailless nucleosomes in Archaea (White and Bell 2002, Koster, Snel et al. 2015). We found that the cleavage of arginine methylated histone tails of +1 nucleosome to generate fragile tailless nucleosome is essential to release the paused Pol II (Liu, Wang et al. 2017, Liu, Wang et al. 2018, Lee, Liu et al. 2020, Liu, Ramachandran et al. 2020).

On the other hand, the late Dr. David Allis found that hyper-acetylation is coupled with transcription activation in vivo (Allis, Glover et al. 1980, Jeppesen and Turner 1993). It was found that neutralized positive charged histone tails via acetylation (Vidali, Gershey et al. 1968) will dramatically compromise the association between DNA and octamer histone subunits from computing (Fenley, Adams et al. 2010). It was reported that hyperacetylation of nucleosomes within the genome leads to easy access by DNase I (Simpson 1978) *in vivo* and fragile *in vitro* (Brower-Toland, Wacker et al. 2005), suggesting “pan-acetylation of nucleosomes” could be a form of fragile nucleosome for Pol II to overcome along gene bodies *in vivo*. *In vitro* transcription assays showed that both pan-acetylated nucleosomes and tailless nucleosomes could confer the ability of T7 RNAP to transcribe, though not as fast as on naked DNA (Protacio, Li et al. 2000). It may be legitimate to claim that both pan-acetylated nucleosomes and tailless nucleosomes are fragile and could be generated along gene bodies for Pol II to transcribe *in vivo*. However, later single-molecule experiments with either a pan-acetylated nucleosome system or a tailless nucleosome system did not show a better advantage of the passage of Pol II compared to the native nucleosome (Bintu, Ishibashi et al. 2012), contradictory to the early reported results for T7 RNAP (Protacio, Li et al. 2000). The results from this report and others force people to look for alternative pathways for Pol II to overcome the resistance of nucleosomes along gene bodies. A common view is that hexasome nucleosomes, which lack an H2A/H2B dimer of an octamer of a regular nucleosome, could be generated by chromatin remodelers with the help of chaperonin complex FACT, and are fragile for Pol II to transcribe over along gene bodies (Rhodes 2024). However, from the report by the late Widom’s group and others, the H3/H4 tetramer seems to contribute most to the stability of nucleosomes (Protacio, Li et al. 2000), suggesting that hexasome nucleosomes are not fragile. Interestingly, it was found that the transcription rate (∼300 nts/min) of Pol II in the artificial assays is much slower than that *in vivo* (>2,000 nts/min) even on naked DNA (Izban and Luse 1992, Bintu, Ishibashi et al. 2012), suggesting artificial *in vitro* assay system does not exactly mimic the action of Pol II *in vivo*.

A recent preprint report showed that the addition of Spt4/Spt5 complex into the artificial assay system dramatically increases the rate of Pol II to ∼2,500 nts/min on naked DNA (Wang 2025), which is similar to that of Pol II *in vivo* while Spt4/5 was missing in the early reports (Izban and Luse 1992, Bintu, Ishibashi et al. 2012). Spt4/5 complex is the only transcription factor conserved from bacteria to human beings, and it is essential for the proper elongation of RNAPs either in bacteria or animals via preventing the backtracking and falling-off of the elongating RNAP from DNA (Kang, Mooney et al. 2018). Without Spt4/5, Pol II is not stable even on DNA alone; any further resistance from tailless nucleosome, pan-acetylated nucleosome, or native nucleosome will aggravate the consequence. There is a need to revisit the early conclusion of the single-molecule experiments regarding tailless and hyperacetylated nucleosomes. However, one of the major missed puzzles in the transcription field is how pan-acetylation of histones along gene bodies is achieved, since acetyltransferase enzymes themselves do not have an engine that would carry them along gene bodies to carry out corresponding modifications. The corresponding PI found that chromatin remodelers act as drivers that allow a variety of histone modification complexes to carry out their functions along gene bodies, including acetyltransferases to generate pan-acetylated nucleosomes along gene bodies (Zhang 2024).

Besides acetylation of histone subunits and cleavage of histone tails, other histone modifications occur, including phosphorylation (Rossetto, Avvakumov et al. 2012, Sawicka and Seiser 2014). The introduction of phosphor-groups in histone tails also weakens the interaction between DNA and histone tails due to the negative charge of phosphor-groups, which could represent another form of fragile nucleosome. However, the exact function of phosphorylation on histone tails in transcription is not well studied. A report showed that phosphorylated H3S10 (H3S10ph) is rich within the gene bodies of actively transcribing genes (Chen, Goyal et al. 2018), suggesting that histone phosphorylation may act, like histone acetylation, to reduce the association between DNA and histone octamers to help Pol II transcribe along gene bodies.

Based on the above analysis, we propose that different forms of fragile nucleosomes are essential for Pol II to transcribe along gene bodies. To prove this hypothesis, here, in using HEK293T cells, we demonstrate that both pan-acetylation and pan-phosphorylation are coupled with transcriptional activation of Pol II. Moreover, we find that all protein-coding genes within HEK293T cells can be classified into four major groups: gene bodies on which pan-acetylation of nucleosomes dominates, gene bodies on which pan-phosphorylation of nucleosomes dominates, gene bodies on which nucleosomes are both acetylated and phosphorylated, and genes that have acetylated nucleosomes at promoters only.

### Pan-acetylation and pan-phosphorylation couple with the activities of Pol II within the entire human genome of HEK293T cells

It is very well known that transcription is coupled with pan-acetylation of histone subunits in eukaryotes (Allis, Glover et al. 1980, Jeppesen and Turner 1993). To understand the underlying mechanism of histone acetylation and Pol II activation in more detail, we carried out **c**leavage **u**nder **t**argets and **tag**mentation sequencing (CUT&TAG-seq) (Skene and Henikoff 2017) analysis of acetylated H3K9 (H3K9ac) in HEK293T cells. We found that H3K9ac, which is generated by the SAGA acetyltransferase complex (Grant, Duggan et al. 1997), was distributed genome-wide (**Fig. 1A**). For comparison, we performed a similar CUT&TAG-seq on acetylated H4K12 (H4K12ac), which is acetylated by the NuA4 acetyltransferase complex (Smith, Eisen et al. 1998). A similar pattern of distribution of H4K12ac to that of H3K9ac was observed (**Fig. 1B**). Since pan-acetylation of histone subunits is coupled with transcription activation (Allis, Glover et al. 1980), we asked if genome-wide distributions of H3K9ac and H4K12ac were coupled with active Pol II (Ser2ph-CTD of Pol II). CUT&TAG-seq of Ser2ph-CTD of Pol II was carried out and a similar genome-wide distribution was found (**Fig. 1C**). To further confirm the authenticity, genome-wide RNA-seq was also carried out; again, the mRNAs are consistent with pan-acetylation of histone subunits and active Pol II distribution genome-wide (**Fig. 1D).**

**Figure 1.**
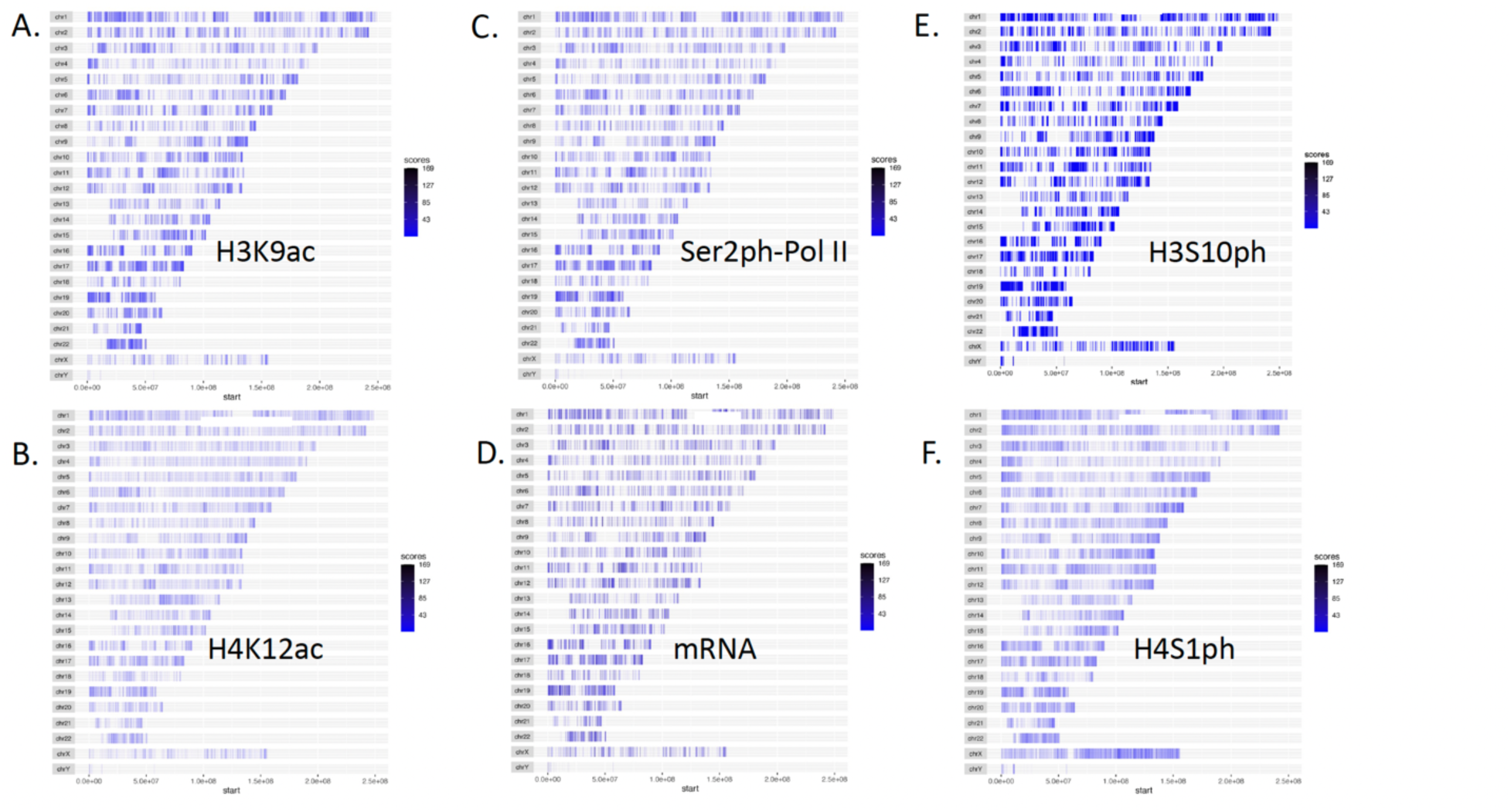
Genome-wide distribution of pan-acetylation, pan-phosphorylation, active Pol II, RNAs in human HEK293T cells. **A**. Acetylated H3K9. **B**. Acetylated H4K12. **C**. Active elongating Pol IIs (Ser2ph-CTD of Pol II). **D**. mRNAs**. E.** Phosphorylated H3S10. **F**. Phosphorylated H4S1.

Histone phosphorylation was well characterized by numerous groups (Hsu, Sun et al. 2000, Zeitlin, Shelby et al. 2001, Sawicka and Seiser 2014, Gil and Vagnarelli 2019). However, it remains one of the missing puzzles whether histone phosphorylation is coupled with transcription activation in eukaryotes, though it was reported that H3S10ph exists within the gene bodies (Chen, Goyal et al. 2018). Following a similar approach for H3K9ac and H4K12ac, we carry out a CUT&TAG-seq for H3S10ph, the most popular histone phosphorylation characterized and generated mostly by MSKs (Mitogen and stress-activated protein kinases) and RSKs (Ribosomal S6 kinases) families (Jin, Wang et al. 1999, Sassone-Corsi, Mizzen et al. 1999, Thomson, Clayton et al. 1999, Wiggin, Soloaga et al. 2002, Soloaga, Thomson et al. 2003). To our big surprise, the distribution of H3S10ph, similar to that of H3K9ac and H4K12ac, is coupled with Ser2ph-CTD of Pol II and mRNAs throughout the entire genome (**Fig. 1E**). This result suggests that the activities of Pol II could also be coupled with the pan-phosphorylation of the nucleosomes within gene bodies. To cross-confirm this speculation, we carried out a CUT&TAG-seq experiment on phosphorylated H4S1 (H4S1ph), another major phosphorylation modification of histone generated by Casein Kinase II (CK2) (Cheung, Turner et al. 2005, Govin, Schug et al. 2010). It shows a similar phosphorylation pattern along the entire genome to that of H3S10ph (**Fig. 1F**). From these data, it suggests that histone phosphorylation has a distribution and function similar to those of acetylation to couple with the activity of Pol II. Moreover, it may be fair to claim that both pan-acetylation and pan-phosphorylation are coupled with transcription of Pol II within gene bodies in human HEK293T cells.

### Genes with pan-acetylation-dominated, pan-phosphorylation-dominated, and both Pan-acetylation and Pan-phosphorylation

Although the results in **Fig. 1** suggest that acetylation, phosphorylation, active Pol II, and mRNAs all overlap, more detailed information emerges when the sites for each of these properties are examined in more detail. We established criteria to categorize the types of modifications that dominated each gene body. Gene bodies that were associated only with acetylated histones were identified by definitive peaks of H3K9ac or H4K12ac and thus grouped as such. Examples of such genes and the sites of their modifications are shown in **Fig. 2A**. We found that approximately 5,009 genes were associated with histones that were dominantly acetylated (**Table 1** and **Table S1**). Genes that had acetylation signals of only ±200 bp from transcription start sites were not included in this group. Phosphorylation-dominated genes were identified by the presence of either H3S10ph or H4S1ph on histones within the gene bodies. Only ∼1,480 genes fell into this group.

**Figure 2.**
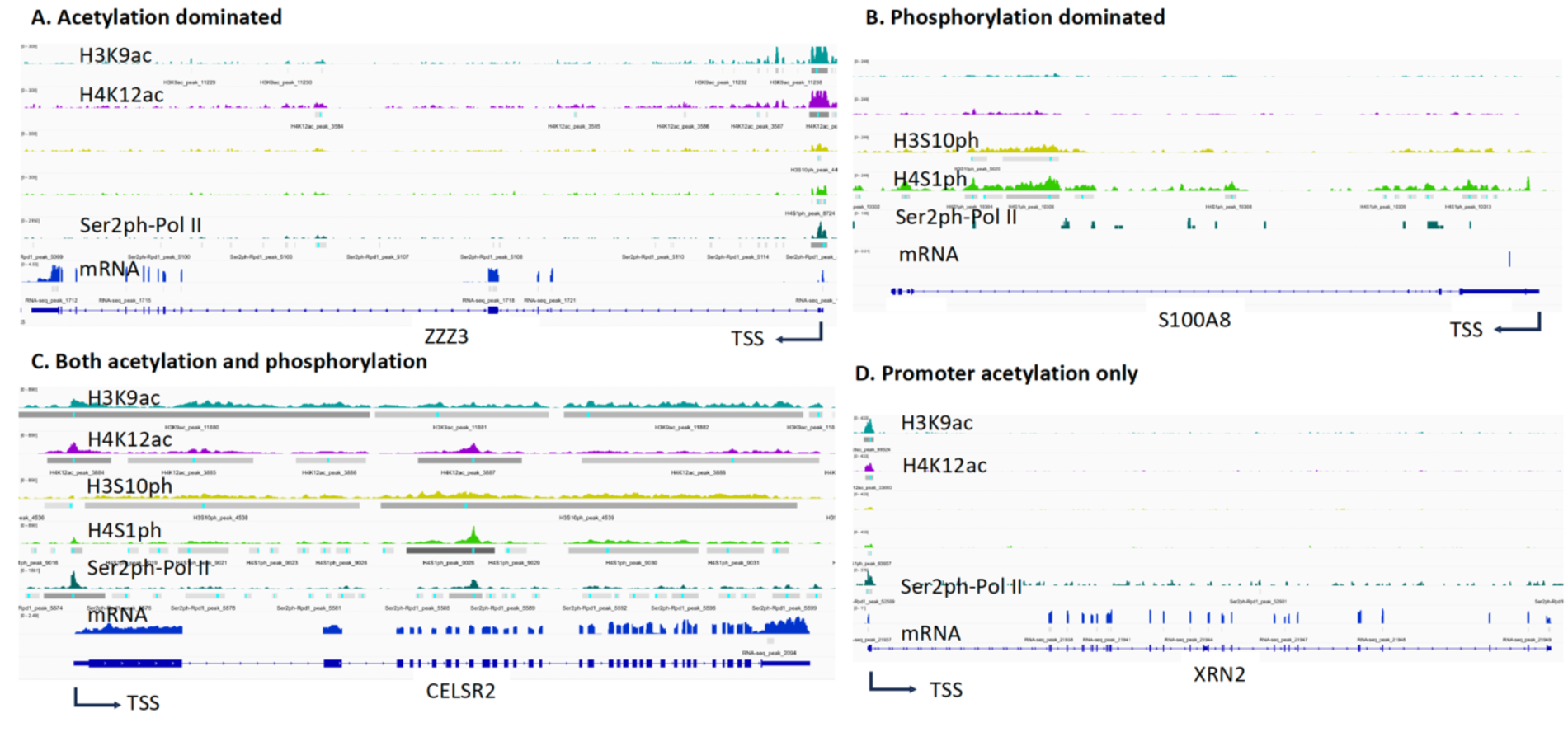
Three types of histone modifications along gene bodies in HEK293T cells and promoter-only acetylation. **A**. Pan-acetylation dominated genes. **B**. Pan-phosphorylation dominated genes. **C**. Both pan-acetylation and pan-phosphorylation genes. **D**. Acetylation at promoters.

**Table 1.** Classified Protein-coding Genes.

|  | Genes |
| --- | --- |
| Acetylation dominated | 5,009 |
| Phosphorylation dominated | 1,480 |
| Both acetylation and phosphorylation | 8,931 |
| Promoter acetylation only | 767 |
| Promoter acetylation and phosphorylation | 3,437 |
| Promoter phosphorylation only | 230 |
|  | 19,854 |

Examples of such genes and the sites of their modifications are shown in **Fig. 2B**) (**Table 1 and Table S2**). Similarly to above, genes that had signals only ±200 bp from their transcription start sites were excluded from this group. Interestingly, most genes (∼8,931) (**Fig. 2C**) (**Table 1** and **Table S3**) were associated with histones that were both acetylated and phosphorylated. Examples of such genes and the types and sites of modifications to their histones are shown in **Fig. 2C**. Again, modifications that were within ±200 bp from their transcription start sites were excluded. Genes on which the only modifications were within ±200 bp of their transcription start sites were placed into a separate set of categories because such modifications may act as docking sites for the recruitment of co-factors during transcription activation, which could remain unchanged after the transcription activation cycle or at a silent status, a potential epigenetic memory for a once-active gene but silent at the status quo. Similar to the criteria of body modification marks described above, these genes were also grouped by the types of modifications observed, acetylation only, phosphorylation only, and both acetylation and phosphorylation (**Fig. 2D**). To our surprise, ∼767 promoters were associated with histones that were only acetylated (**Table 1 and Table S4**), ∼230 promoters had histones that were only phosphorylated (**Table 1 and Table S5**), while most of these promoters, ∼3,437, were associated with histones that were acetylated and phosphorylated (**Table 1 and Table S6**). From these data, we speculate that phosphorylation of histones at promoter regions, especially -1 nucleosome, may not exist. The signals we derived could be caused by noise, though an early report showed phosphorylation of histone subunits at the promoter and enhancer-regulated genes (Kellner, Ramos et al. 2012). It is likely that acetylation of histone subunits at -1 nucleosomes could be a major modification as docking sites to recruit other transcription co-factors and last permanently even after a transcription cycle (Zhang 2024). Since the current ChIP-seq and CUT&TAG-seq technologies could not discriminate +1 or -1 nucleosomes, our data suggest there exist acetylation modifications at -1 nucleosomes, which remain to exist even after a transcription activation cycle. Conversely, all modifications within gene bodies (including +1 nucleosome) should be dynamic and transient. If we add all genes classified above together with a total of ∼19, 854 genes (**Table 1**) out of the human HEK293T genome, with a total of 19,989 protein-coding genes (GRCh38). We assume that the small number of left genes (∼135) without any modifications could be silent within human HEK293T cells, but could be triggered in other differentiated cell types since HEK293T cells were immortalized human embryonic kidney cells. Another highly likely possibility is that the modification signals are too weak to be detected due to systemic errors. One bold or provocative guess is that all genes expressed by Pol II within HEK293T cells, either active or inactive, have epigenetic marks.

### Inhibition of Pol II by DRB prevents transcription initiation of elongation and shuts down all transcription

To prove that both pan-acetylation and pan-phosphorylation of nucleosomes along gene bodies are essential for the elongation of Pol II, we need to establish a readout assay to show the coupling of both modifications and active Pol II. DRB(5,6-dichloro-1-beta-d-ribofuranosylbenzimidazole) was found to inhibit CDK7 (Yankulov, Yamashita et al. 1995), CDK9 (Baumli, Endicott et al. 2010), and CK2 (Zandomeni, Zandomeni et al. 1986). A report showed that inhibition of murine myoblasts by DRB could lead to the disappearance of newly synthesized RNAs for all transcripts in 30 min in murine myoblasts (Hou, Wang et al. 2019). The exact underlying working mechanism of DRB on the transcription of Pol II still remains ambiguous. To investigate the effect of DRB on Pol II in human cells, we added DRB to HEK293T cells for one hour and examined the phosphorylation status of Ser5-CTD of Pol II (Ser5ph-CTD of Pol II). To our surprise, Ser5ph-CTD of Pol II is almost completely gone after inhibition of DRB for one hour (**Fig. 3A**) while neither inhibitor of CDK7 nor CDK8 alone does not make any changes to the level of Ser5ph-CTD of Pol II. This result may explain why inhibition of DRB leads to the shutdown of all transcription activities in murine myoblasts (Hou, Wang et al. 2019), a complete shutdown of the initiation (phosphorylation of Ser5-CTD of Pol II) of all Pol II-dependent transcription units. To further prove the notion, we carried out a PRO-seq analysis of human HEK293T cells under inhibition of DRB with time course (**Fig. 3B**). The transcription of all genes started to shut down at 5 min, and the transcription for all genes gradually disappeared after 15 min and 30 min later (**Fig. 3B**). Inspecting individual genes, we do see the gradual disappearance of newly synthesized RNAs (**Fig. 3C**). The results suggest that DRB inhibits the initiation of elongation of Pol II to shut down transcription of Pol II-dependent genes in human HEK293T cells. The working mechanism of DRB inhibition is finally revealed. More importantly, we have established a readout assay on the shutdown of all active Pol IIs *in vivo*.

**Figure 3.**
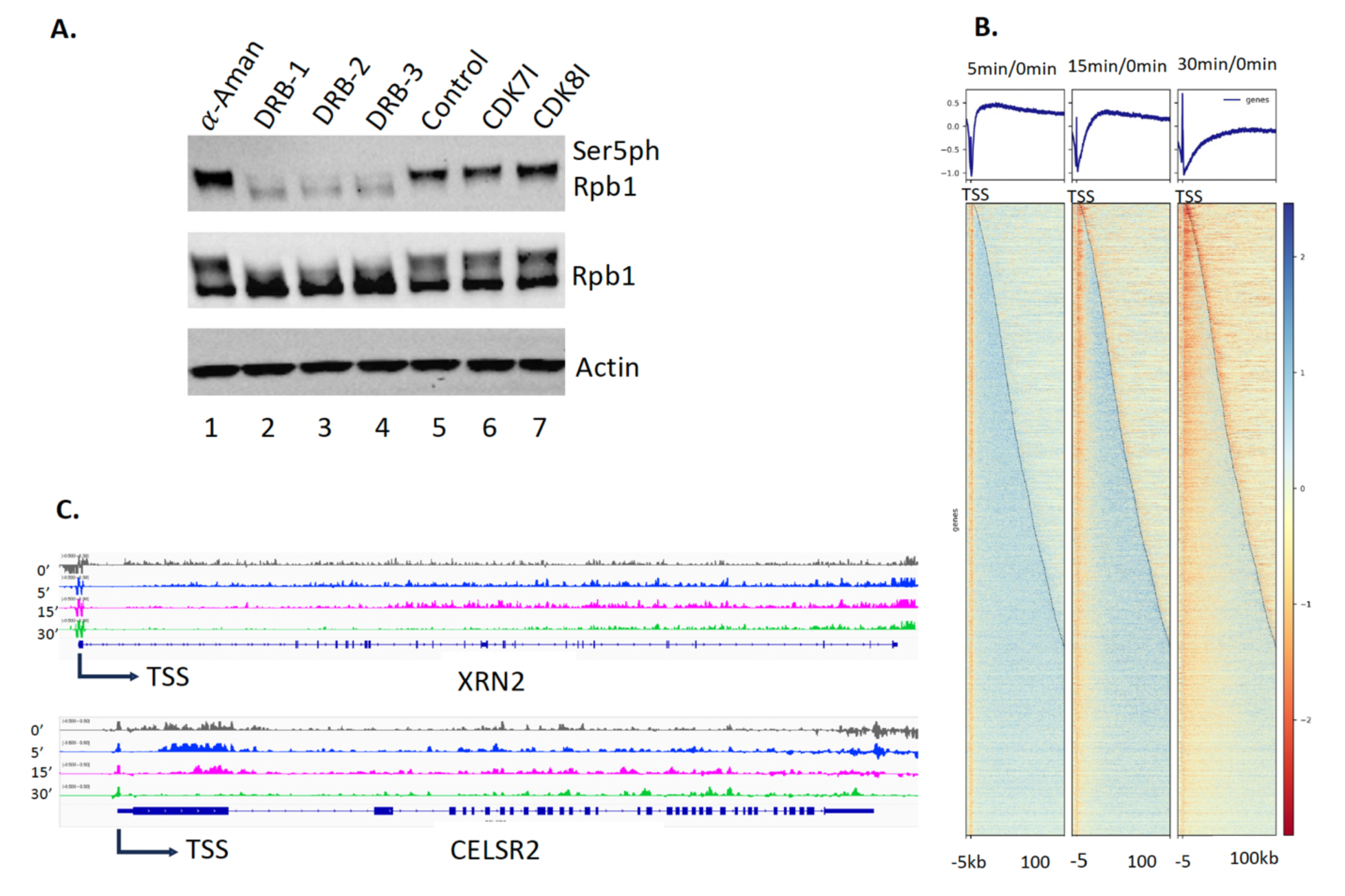
Transcription is completely shut down by DRB (for a total of 4,772 no-overlapping genes). **A**. DRB completely inhibits the phosphorylation of Ser5-CTD of Rpd1 of Pol II. **B.** PRO-seq results of genome-wide of HEK293 after inhibition by DRB at 5 minutes, 15 minutes, and 30 minutes. **C.** Examples of the gradual shutdown of genes with time course after adding DRB.

### The activities of Pol IIs resume with the removal of DRB

After the readout of the shutdown of all active Pol IIs, we still need an assay to read out the activation of the entire genome. It was reported that Triptolide could not only completely shut down the phosphorylation of Ser5-CTD of Pol II through covalently binding to the ATPase protein XPB, a subunit of TFIIH, but also led to the degradation of Pol II (Titov, Gilman et al. 2011). However, the inhibition of DRB does not trigger the degradation of Pol II but leads to the disappearance of both Ser5ph-CTD of Pol II (**Fig. 3A**) and Ser2ph-CTD of Pol II (**Fig. 4A**). Similar to the results of PRO-seq shown above, CUT&TAG-seq of active Pol IIs (Ser2ph-CTD of Pol II showed a wave of the drop of active Pol II along gene bodies though not as sharp as that of PRO-seq (**Fig. 4B**). We reason that the removal of DRB in the cell growing medium should resume the activity of all Pol IIs, which are just shut down by DRB. After inhibition of DRB for 30 minutes, the medium of HEK293T cells was replaced with a DRB-free medium, and a CUT&TAG-seq of active Pol IIs (Ser2ph-CTD of Pol II detected by 3E10 antibody) with 10 minutes, 30 minutes, and 60 minutes recovery time was carried out. To our satisfaction, active Pol IIs come back along gene bodies (**Fig. 4C**). To our surprise, CUT&TAG-seq of active Pol IIs could precisely read out the activation process of Pol IIs along gene bodies (**Fig. 4C).**

**Figure 4.**
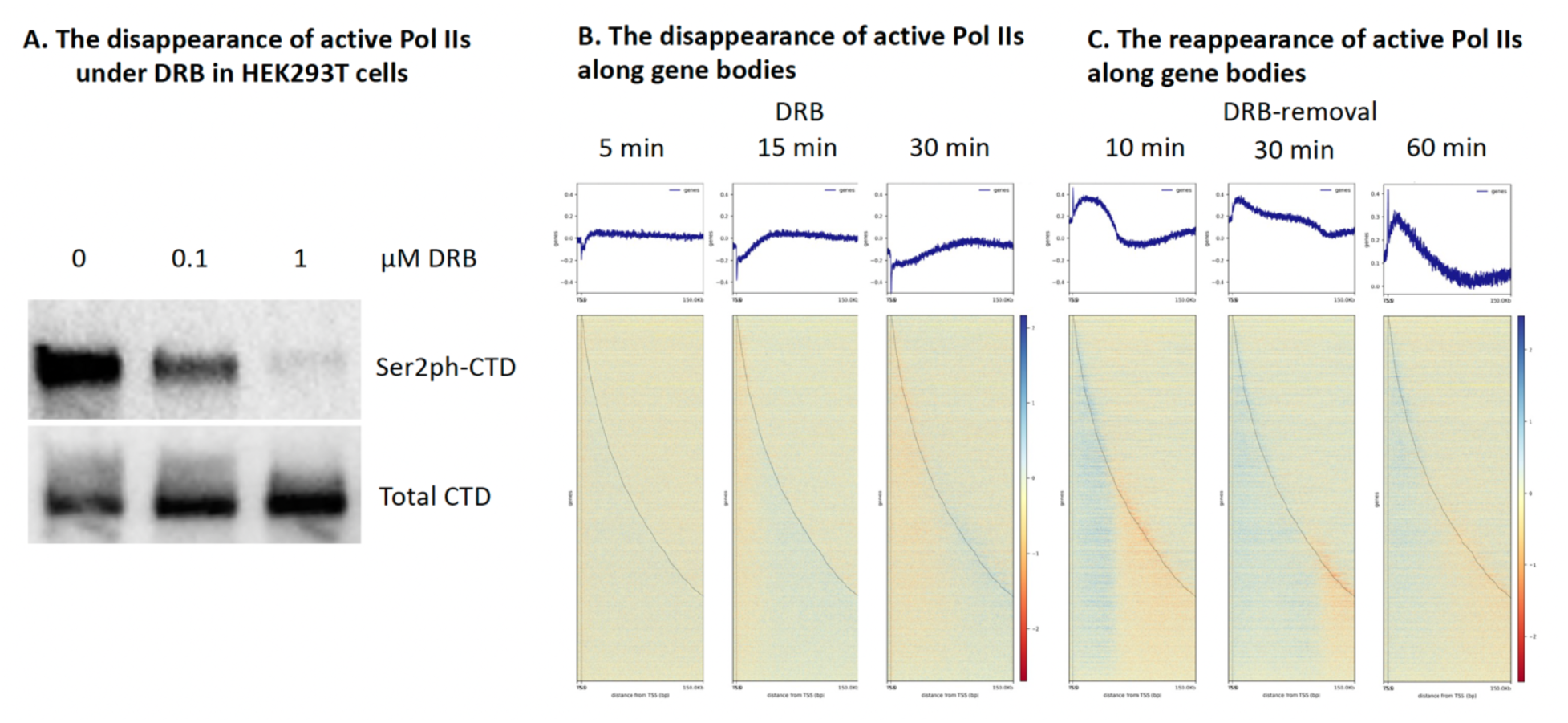
Active Pol IIs are lost with the addition of DRB while they come back with the removal of DRB (for a total of 4,772 no-overlapping genes). **A**. Active Pol IIs gradually disappear with DRB. **B**. Active Pol IIs come back with the removal of DRB.

### Transcription activation of Pol IIs is coupled with pan-acetylation along gene bodies

It was well established that pan-acetylation is coupled with transcription activation of Pol II (Allis, Glover et al. 1980, Jeppesen and Turner 1993). The PI proposed that all nucleosomes along gene bodies should be fully acetylated for these groups of genes, but the modifications should be transient due to the action of Pol II and subsequent rebuilding of nucleosomes after the leave of elongating Pol IIs (Zhang 2024). Our data showed that active Pol II, mRNA, H3K9ac, and H4K12ac are coupled (**Fig. 1A, 1B, 1C, 1D**), which further consolidates the coupling fact of the activities of Pol II and pan-acetylation of gene bodies. However, if the activities of Pol II are coupled with pan-acetylation as we proposed, we should detect the changes of this modification with and without inhibition of DRB. We reason that pan-acetylation should drop or be downregulated when the activity of Pol II is inhibited by DRB while it should come back when the inhibition is removed. This turns out to be true, we saw a drastic drop in the level of H3K9ac overall after the inhibition of DRB (**Fig. 5A**). We carried out a similar CUT&TAG-seq on H3K9ac under the inhibition of DRB for 5 min, 15 min, and 30 min and recovery with a time course of 10 min, 30 min, and 60 min. As predicted, with the inhibition of DRB, the level of H3K9ac dropped while it came back when DRB was removed (**Fig. 5A**). This is also true for H4K12ac (**Fig. 5B and Fig. S1**). The results prove the notion that pan-acetylation is coupled with the activation of Pol II. From these results, inhibition of DRB led to a drop in the level of H3K9ac and H4K12ac. In our repeated experiments, we saw a gradual drop more obviously in the level of both H3K9ac and H4K12 ac as well (**Fig. S2**). As classified previously, there are three major types of histone modifications, we only included 5,009 acetylation-dominated genes, plus 8,931 genes with both acetylation and phosphorylation, with a total of 13,940 genes for this final analysis.

**Figure 5.**
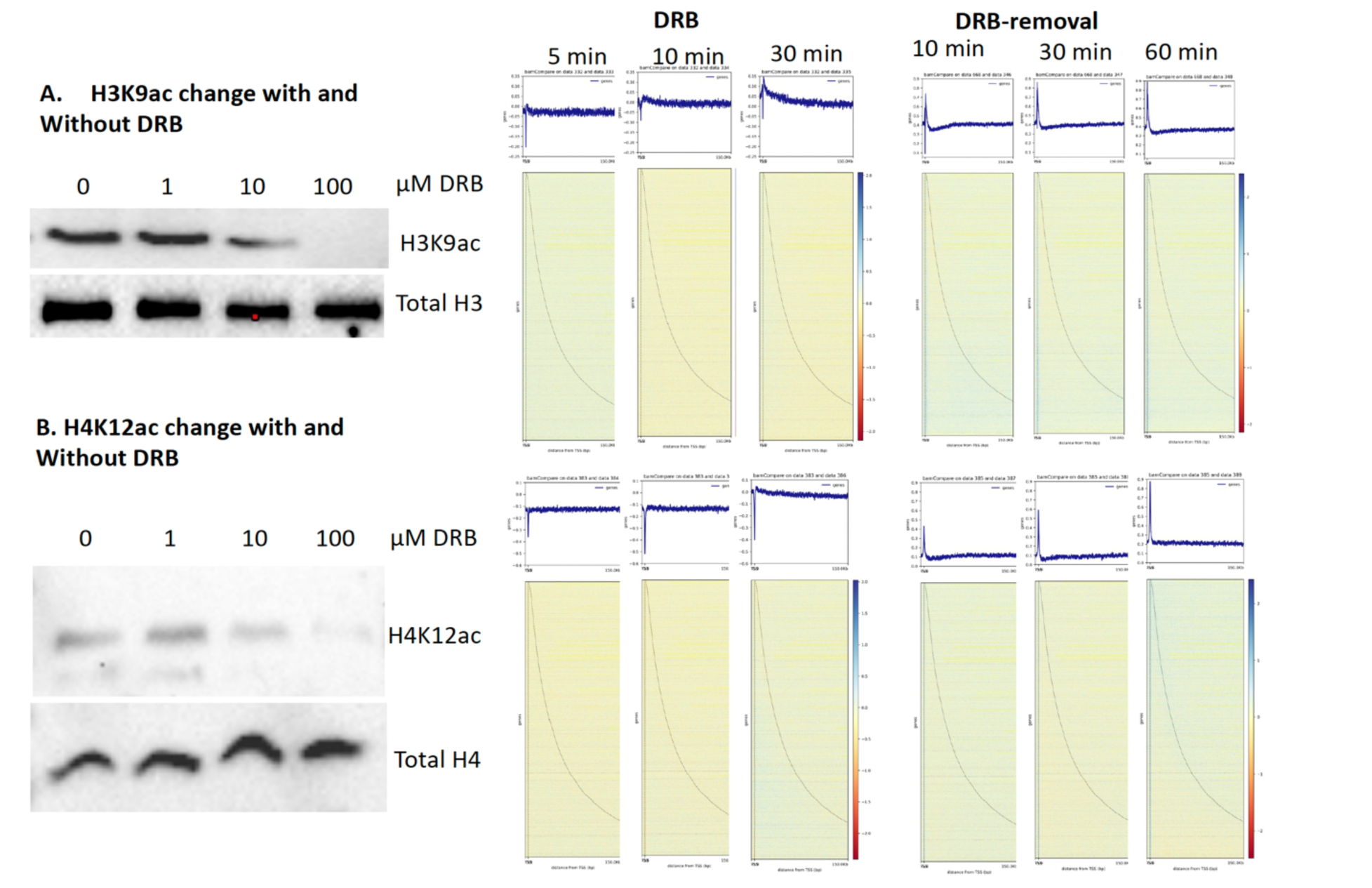
Pan-acetylation couples with transcription activation (total of 13,940 genes, with 5,009 acetylation-dominated genes and with 8,931 both acetylation and phosphorylation). **A.** DRB inhibition leads to the drop of H3K9ac while H3K9ac comes back with the removal of DRB. **B.** The change of H4K12ac has a similar pattern to that of H4K12ac.

### Transcription activation of Pol II is coupled with pan-phosphorylation along gene bodies

We find that pan-phosphorylation is also coupled with activities of Pol II (**Fig. 1**). It should be no surprise that similar effects of H3S10ph and H4S1ph to those of H3K9ac and H4K12ac should be observed after inhibition of Pol II by DRB and come back after removal of DRB. It turns out to be true that inhibition of DRB led to a significant drop of H3S10ph (**Fig. 6A**, **Fig. S3**). H3S10ph and H4S1ph along gene bodies are dropped after inhibition of DRB at 5 min, 15 min, and 30 min, respectively, while both modifications came back when DRB was removed after 10 min, 30 min, and 60 min (**Fig. 6**). In our repeating experiments, the drop of both modifications also showed different dynamics (**Fig. S4**). Overall, this result further supports our speculation that pan-phosphorylation is as important as pan-acetylation to couple the activities of Pol II by generating “fragile nucleosomes” along gene bodies for Pol II to transcribe. Since most genes are regulated by both pan-acetylation and pan-phosphorylation, it was proposed that pan-acetylation could happen in the pioneering run of transcription (Zhang 2024). After the first run, there exist two possibilities: one continues to be regulated by pan-acetylation (Zhang 2024), while another one could be regulated by phosphorylation. It was reported that acetylation is ahead of phosphorylation, while acetylation does not affect phosphorylation, but phosphorylation prevents acetylation (Mahadevan, Willis et al. 1991, Cheung, Tanner et al. 2000). These results suggest that pan-acetylation may happen ahead of pan-phosphorylation.

**Figure 6.**
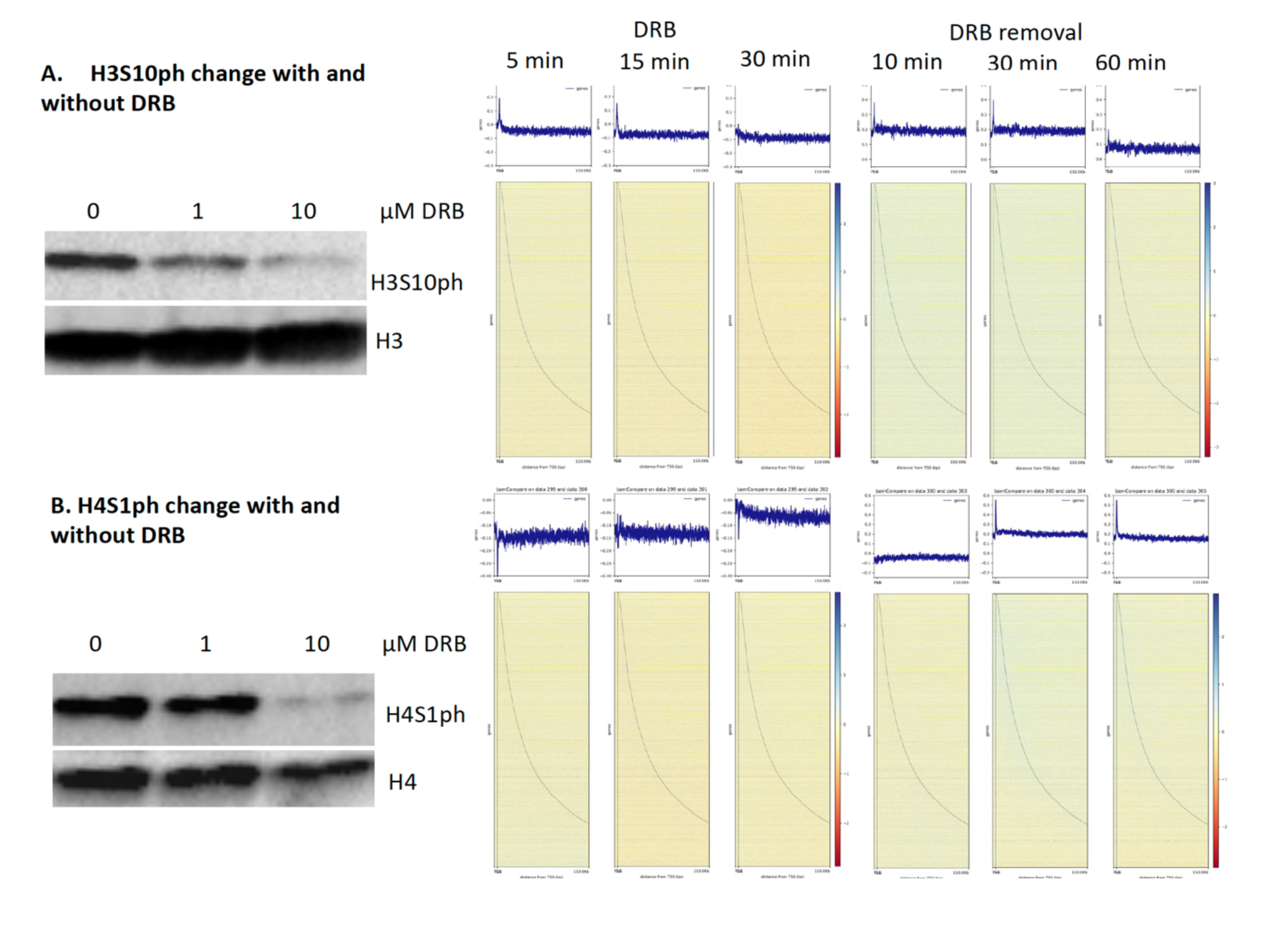
Pan-phosphorylation couples with transcription activation (Total of 10,411 genes, with 1480 phosphorylation dominated and with 8,931 both acetylation and phosphorylation). **A.** DRB inhibition leads to the drop of H3S10ph while H3S10ph comes back with the removal of DRB. **B.** The change of H4S1ph has a similar pattern to that of H3S10ph.

One possible explanation is that pan-acetylation happens at the first round or pioneer round of transcription of Pol II, then pan-phosphorylation follows in the second round and later rounds. Again, the analysis of the above is focused on pan-phosphorylation-dominated genes (1,480) plus both pan-acetylated and pan-phosphorylated genes (8,931) with a total of 10,411 genes.

## DISCUSSION

Nucleosomes are barriers and prevent Pol II from transcribing along gene bodies in eukaryotes (Kornberg and Thomas 1974, Olins and Olins 1974, Kireeva, Hancock et al. 2005). It has become a long-standing but not well-appreciated puzzle in the field. There are numerous models to interpret how Pol II overcomes the resistance of nucleosomes *in* vivo (Aoi and Shilatifard 2023). However, the exact underlying working mechanism still remains ambiguous. A preferred model or argument is that Pol II will transcribe on “hexasome nucleosomes,” which lack an H2A/H2B dimer of octamer nucleosome. It was claimed that hexasome nucleosomes are fragile (Baer and Rhodes 1983, Weiner, Hughes et al. 2010, Henikoff, Belsky et al. 2011, Xi, Yao et al. 2011, Kubik, Bruzzone et al. 2015) and could be generated during Pol II elongation (Rhee, Bataille et al. 2014, Ramachandran, Ahmad et al. 2017). It was reported that these hexasomes are generated by chromatin remodelers (Brahma and Henikoff 2019, Klein, Troy et al. 2023, Wu, Munoz et al. 2023, Zhang, Jungblut et al. 2023, Rhodes 2024) with the help of chaperone complex FACT (Zhou, Liu et al. 2020). Nevertheless, an early sophistical in vitro experiment carried out by the late Dr. Jonathan Widom’s group showed that pan-acetylated nucleosomes derived from Hela cells and artificial “tailless nucleosomes” confer the ability of T7 RNAP to transcribe on these nucleosomes ∼10-20 times faster than on a native nucleosome (just one nucleosome system) without any modifications (Protacio, Li et al. 2000). Most importantly, the experiment showed that the tails of H2A/H2B had a minor effect on the speed of T7 RNAP, while tails of H3/H4 played a dominant role in blocking the pass of T7 RNAP (Protacio, Li et al. 2000). The result suggests that the hexasome nucleosome with intact H3/H4 is not fragile, which still could block RNAPs or Pol II transcription along gene bodies. In this regard, how Pol II overcomes the resistance of nucleosomes during transcription becomes the most urgent question in the field, though it has been missed completely in the past two decades.

We serendipitously revealed that arginine methylated histone tails on +1 nucleosomes for a large group of genes are cleaved to generate “tailless nucleosome” for Pol II to release from a paused status in metazoans (Liu, Wang et al. 2017, Liu, Wang et al. 2018, Lee, Liu et al. 2020, Liu, Ramachandran et al. 2020, Liu, Wei et al. 2022). The novel discoveries remind us of the existence of the “tailless nucleosome” in Archaea bacteria (White and Bell 2002, Koster, Snel et al. 2015), of which RNAP could transcribe the entire genome without the assistance of any epigenetic modifications or the help of chromatin remodelers, which is always the case in eukaryotes. Based on the property of “tailless nucleosome” found in metazoans, in Archaea bacteria, and in vitro transcription assays, it is safe to claim that “tailless nucleosome” is one of the forms of “fragile nucleosomes,” which is essential to be generated to release paused Pol II in metazoans (Zhang 2024) (**Fig. 7**).

**Figure 7.**
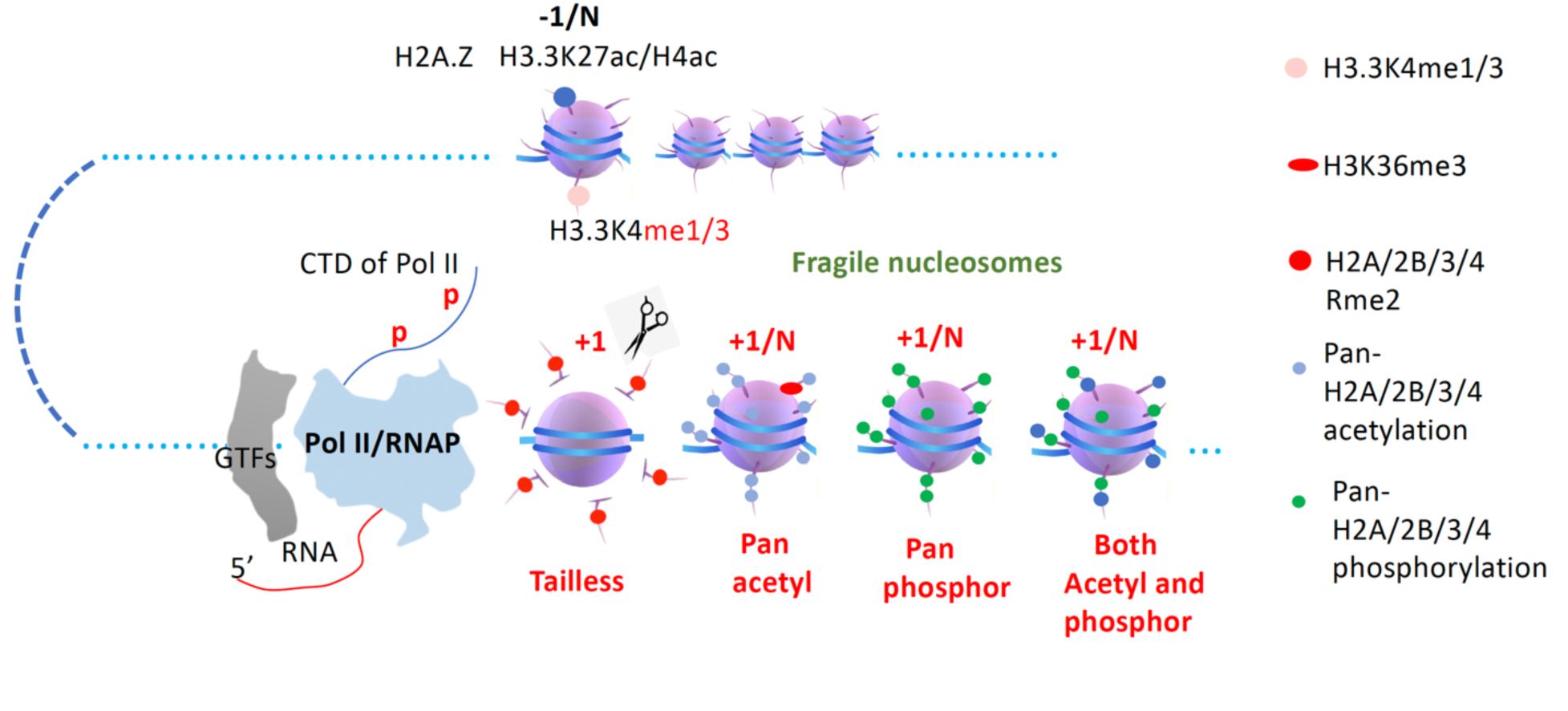
A model of the general transcription regulation mechanism of Pol II in eukaryotes. Methylated H3.3K4 usually exists at the promoters (-1) and the enhancers (-N). Four forms of fragile nucleosomes, including “tailless,” “pan-acetylated,” “pan-phosphorylated,” and “both acetylated and phosphorylated” within gene bodies (+1/N), are generated in metazoans for Pol II to transcribe along gene bodies smoothly.

Pan-acetylated nucleosomes also lead to the compromised interaction between DNA and histone octamer (Simpson 1978, Brower-Toland, Wacker et al. 2005). Most interestingly, pan-acetylated nucleosomes behave similarly to that of tailless nucleosomes to confer the ability of RNAP or Pol II to transcribe in vitro (Protacio, Li et al. 2000). This is consistent with the discovery of the late Dr. David Allis, pan-acetylation is always coupled with transcription activation of Pol II in vivo (Allis, Glover et al. 1980). Individual acetyltransferase and deacetylase have been intensively studied in the field in recent decades since their early discoveries (Brownell, Zhou et al. 1996, De Rubertis, Kadosh et al. 1996). However, people remain puzzled about how pan-acetylation and deacetylation along gene bodies are involved in the activities of Pol II during a transcription activation cycle in vivo, though two main functions have been assigned (Winston and Allis 1999, Strahl and Allis 2000, Roth, Denu et al. 2001). When the driver function of chromatin remodelers was assigned, pan-acetylation of nucleosomes along gene bodies could be achieved *in vivo* (Zhang 2024). Here, we confirmed that “Pan-acetylated nucleosomes” are ubiquitous along gene bodies and coupled with activities of Pol II in HEK293T cells. It is likely that “pan-acetylated nucleosome” is one of the most popular forms of “fragile nucleosomes” in eukaryotes (**Fig. 7**).

Most surprisingly, the role of pan-phosphorylation along gene bodies to assist Pol II is not well appreciated or completely ignored in the field so far. Furthermore, it is still an unexplored area on how the pan-phosphorylation along gene bodies is deposited, though numerous histone kinase candidates that have been well characterized (Rossetto, Avvakumov et al. 2012). Based on our serendipitous discoveries of the pan-phosphorylation coupling with the activity of Pol II, it may be safe to claim that pan-phosphorylation of nucleosomes is as important as that of pan-acetylation of nucleosomes to generate “fragile nucleosomes” along gene bodies to help Pol II overcome the resistance of tightly associated DNA and octamer of nucleosome during transcription elongation in all eukaryotes (**Fig. 7**). Furthermore, there also exist nucleosomes with mixed acetylation and phosphorylation within the bodies of a majority group of genes in eukaryotes, which also belong to the “fragile nucleosomes” for Pol II to transcribe smoothly along gene bodies (**Fig. 7**).

In conclusion, the ultimate goal of all epigenetic modifications in eukaryotes is to serve the activities of Pol II, which could be manifested by the generation and degeneration of fragile nucleosomes along gene bodies. Overall, there are at least three forms of fragile nucleosomes in yeast and four forms of fragile nucleosomes in animals. Pol II could transcribe along all four fragile nucleosomes smoothly in eukaryotes. Combined with the transcription of RNAPs from bacteria and Archaea, a general model of transcription of all RNAPs from bacteria to human beings could be derived (**Fig. 8**).

**Figure. 8.**
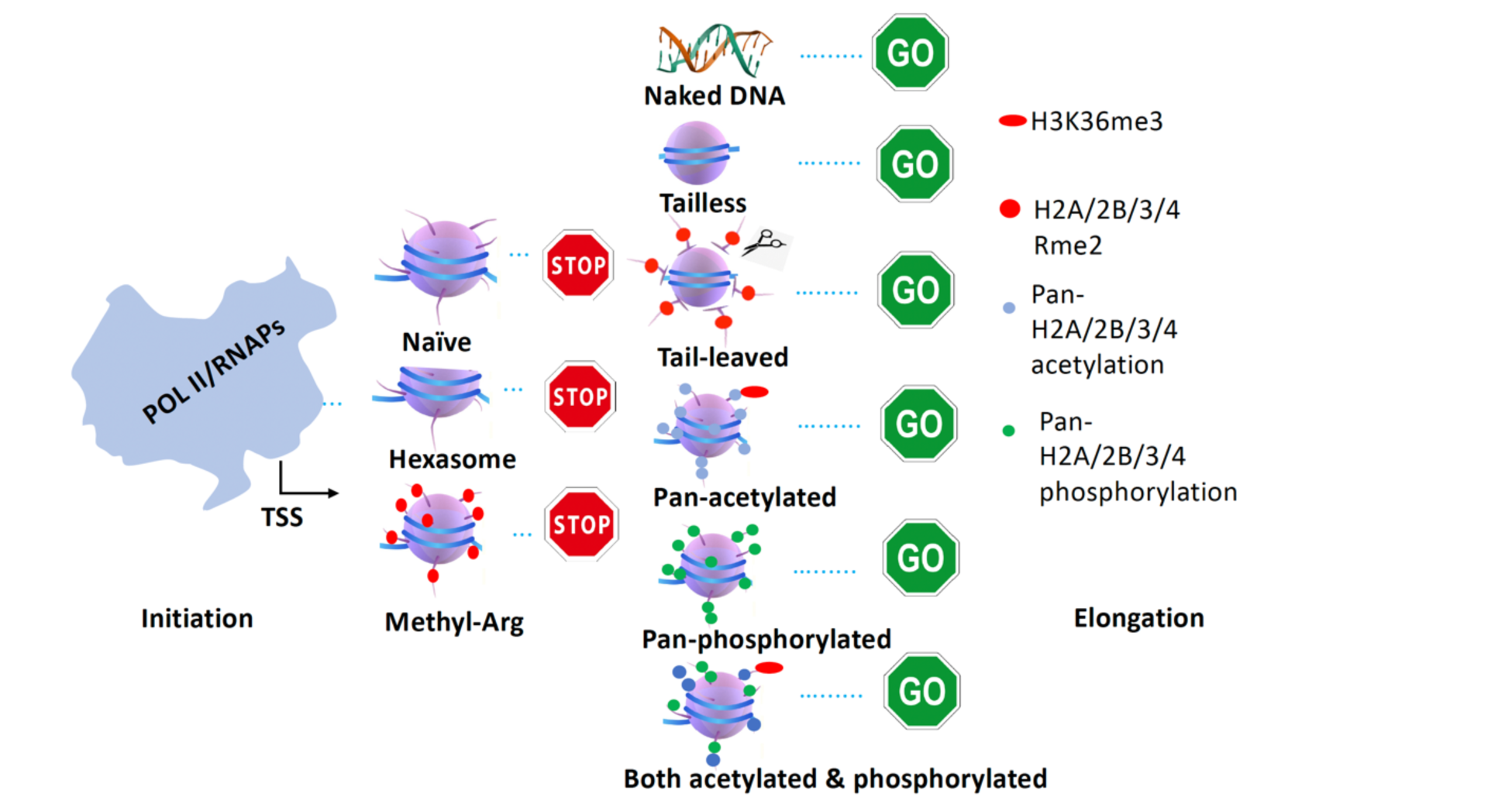
A general model of transcription from bacteria to human beings. RNAPs could be blocked by nucleosomes or hexasomes with or without arginine methylation. RNAPs could transcribe on naked DNA (in bacteria), tailless nucleosomes (in Archaea), tail-cleaved nucleosomes (+1 nucleosome for some genes in metazoans), pan-acetylated nucleosomes, pan-phosphorylated nucleosomes, both pan-acetylated and pan-phosphorylated nucleosomes.

## Materials and Methods

### CUT&TAG-seq

CUT&TAG-seq experiments were carried out following the protocol by Henikoff v3 (https://www.protocols.io/) with some minor changes. **Preparing ConA beads**. Gently resuspend and withdraw enough of the ConA bead slurry such that there will be 10 μL for each final sample of up to 500,000 cells. The following is for all samples. Transfer 170 µL ConA bead slurry into 1.6 mL Binding buffer in a 2 mL tube and mix by pipetting. Place the tube on a magnet stand to clear (30 s to 2 min). Withdraw the liquid completely and remove it from the magnet stand. Add 1.5 mL Binding buffer, mix by pipetting, and remove liquid from the cap with a quick pulse on a microcentrifuge. Place on a magnet stand to clear, withdraw liquid, and resuspend in 170 µL Binding buffer (10 μL per sample) and hold the bead slurry at room temperature until cells are ready. **Prepare lightly fixed cells**. HEK293T cells were cultured in DMEM supplemented w/10%FCS. Split cells into 1 million cells/ per 10cm dish 1 day before harvesting. Wash cells 2 times in PBS. Add 4 ml 0.1% 16% formaldehyde in PBS into each dish. and incubate at room temperature for 2 minutes. Stop cross-linking by addition of 1.25 M glycine to twice the molar concentration of formaldehyde (e.g., 600 µL to 10 ml). Wash 2 times in PBS and harvest cells in PBS in conical centrifuge tubes (15 ml or 50 ml). Centrifuge **4 min** 1300 x g at 4 °C and drain liquid by pouring off and inverting onto a paper towel for a few seconds. Resuspend in Wash buffer to a concentration of ∼1 million cells per ml. Check nuclei using ViCell or cell counter slide. Resuspend in at least 1 volume Wash buffer at room temperature, centrifuge 3 min 600 x g at room temperature, and withdraw liquid. Resuspend in 1.5 mL Wash buffer, and transfer to a 2 mL tube, and hold on ice. **Bind cells to Concanavalin A-coated beads**. Vortexing beads gently and slowly add the bead slurry into the cells. There will be 10 μL for each final sample of up to 500,000 cells. Place on an end-over-end rotator for 5-10 min. **Binding primary antibody**. After a quick spin to remove liquid from the cap (<100 x g), place the tubes on a magnet stand to clear and withdraw the liquid. Resuspend cells in 800 µL ice-cold Antibody buffer with gentle vortexing, place on ice, and divide into 16 1.5 mL tubes, 50 µL each (for 16 samples; scale up or down as needed). Add 0.5-1 µL primary antibody to each sample with gentle vortexing. Nutate (or rotate) at room temperature overnight at 4 °C. Liquid should remain in the bottom and on the side of the tube while rocking. **Binding secondary antibody.** After a quick spin to remove liquid from the cap (<100 x g), place each tube on the magnet stand to clear and pull off the liquid. Mix the secondary antibody 1:100 in Dig-wash buffer and squirt in 100 µL per sample while gently vortexing to allow the solution to dislodge the beads from the sides. Place the tubes on a nutator at room temperature for 30–60 min.00:30:00. After a quick spin, place the tubes on a magnet stand to clear and withdraw the liquid. Add 1 mL Dig-wash buffer. Invert 10x or gently vortex to allow the solution to dislodge most or all of the beads. Repeat the last two steps twice. **Bind pA-Tn5 adapter complex.** Mix pA-Tn5 adapter complex in Dig-300 buffer to a final concentration of 1:250 for 100 µL per sample. After a quick spin, place the tubes on the magnet stand to clear and pull off the liquid. Squirt in 100 µL of the pA-Tn5 mix while gently vortexing to allow the solution to dislodge most or all of the beads. Place the tubes on a nutator at room temperature for 1 hr.01:00:00. After a quick spin, place the tubes on a magnet stand to clear and pull off the liquid. Add 1 mL Dig-300 buffer. Invert 10x or gently vortex to allow the solution to dislodge most or all of the beads. Repeat the last two steps twice. **Tagmentation**. After a quick spin, place the tube on the magnet stand to clear and pull off the liquid. Add 300 µL Tagmentation buffer while gently vortexing. Incubate at 37 °C for 1 hr in a water bath or incubator.01:00:00. **DNA extraction**. To stop tagmentation and solubilize DNA fragments, add 10 µL 0.5M EDTA, 3 µL 10% SDS, and 2.5 µL 20 mg/mL Proteinase K to each sample. Mix by full-speed vortexing ∼2 s, and incubate 1 hr 55 °C to digest (and reverse cross-links). Add 300 μL PCI and mix by full-speed vortexing ∼2 s. Transfer to a phase-lock tube, and centrifuge for 3 min at room temperature at 16,000 x g. Add 300 μL Chloroform and invert ∼10x to mix (do not vortex). Centrifuge 3 min room temperature at 16,000 x g. Remove the aqueous layer by pipetting to a fresh 1.5 mL tube containing 750 µL 100% ethanol, pipetting up and down to mix. Chill on ice and centrifuge for at least 10 min at 4 °C 16,000 x g. Carefully pour off the liquid and drain on a paper towel. There is typically no visible pellet. Rinse in 1 mL 100% ethanol and centrifuge for 1 min at 4 °C 16,000 x g. Carefully pour off the liquid and drain on a paper towel. Air dry. When the tube is dry, dissolve in 25-30 μL 1 mM Tris-HCl pH 8, 0.1 mM EDTA, and vortex on full to dissolve. **PCR**. 21 µL DNA + 2 µL of 10 µM Universal or barcoded i5 primer + 2 µL of 10 µM uniquely barcoded i7 primer. In a thin-wall 0.5 ml PCR tube, using a different barcode for each sample (Indexed primers described by Buenrostro, J.D. et al. Single-cell chromatin accessibility reveals principles of regulatory variation. Nature 523:486 (2015)). Add 25 µL NEBNext HiFi 2x PCR Master mix. Mix, quickly spin, and place in Thermocycler and begin cycling program with heated lid. Cycle 1: 72 °C for 5 min (gap filling), Cycle 2: 98 °C for 30 sec, Cycle 3: 98 °C for 10 sec, Cycle 4: 63 °C for 10 sec, Repeat Cycles 3-4 13 times, 72°C for 1 min and hold at 8 °C. **Post-PCR clear-up**. After the tubes have been cooled, remove them from the cycler and add 1.3 volume (65 µL) SPRI bead slurry, mixing by pipetting up and down. Quick spin and let sit at room temperature for 5-10 min. Place on magnet and allow to clear before carefully withdrawing liquid. On the magnet and without disturbing the beads, add 200 µL 80% ethanol. Withdraw liquid with a pipette to the bottom of the tube and add 200 µL 80% ethanol. Withdraw the liquid, and after a quick spin, remove the remaining liquid with a 20 µL pipette. Proceed immediately to the next step. Remove from magnet stand, add 25 µL 10 mM Tris-HCl pH 8, and vortex on full. After 5 min, place on a magnet stand and allow to clear. 00:05:00. Remove liquid to a fresh 0.5 ml tube with a pipette. **DNA sequencing and data processing**. Determine the size distribution and concentration of libraries by *capillary electrophoresis* using an Agilent 4200 TapeStation with D1000 reagents or equivalent. Libraries were pooled and sequenced on Illumina HiSeq2000 or Illumina NextSeq500 (University of Colorado Anschutz Medical campus Genomics core facility). Data were analyzed by workflows on Basepair (https://www.basepairtech.com).

### RNA-seq

*HEK293T* cell preparation is similar to that of CUR&TAG-seq experiments. For RNA, total RNA was purified with an RNeasy Mini kit (QIAGEN). mRNAs were isolated using NEBNext Poly(A) mRNA Magnetic isolation kit (NEW ENGLAND Biolabs). Libraries were prepared by using NEBNext® Ultra™ RNA Library Prep Kit for Illumina® NEB #E7530S/L (NEW ENGLAND Biolabs). The DNA library was validated using TapeStation (Agilent Technologies). Libraries were pooled and sequenced on NovaSEQ6000 (paired End 2x150bp). Data were analyzed by workflows on Basepair (https://www.basepairtech.com).

### CUT&TAG-seq data process and Figure 1 generation

In this study, we performed an in-depth analysis of chromatin modifications, active Pol II binding, and mRNA production leveraging the CUT&TAG-seq sequencing technology. The obtained raw sequence reads were aligned to the reference genome utilizing the Burrows-Wheeler Aligner (BWA) tool. After the alignment process, we applied the MACS2 (Model-based Analysis of ChIP-seq) peak-calling algorithms to identify regions of significant enrichment. These regions suggest modified chromatin states and potential active Pol II binding. To visualize these areas of chromatin modifications and Pol II binding across various experimental conditions, we utilized the ggplot2 package in R to generate Figure 1. All CUT&TAG-seq and RNA-seq raw data were processed by Basepair (https://app.basepairtech.com/). Bam files from Basepair were used in DeepTools of Galaxy (https://usegalaxy.eu/) to generate bamCompare files, computeMatricx files, and final heatmap figures (Ramirez, Ryan et al. 2016).

### CUT&TAG-seq analysis of H3K9ac, H4K12ac, H3S10ph, H4S1ph, Ser2ph-CTD of Pol II, and RNA-seq

Adapter sequences from reads were trimmed with fastp v0.20.0 (Chen, Zhou et al. 2018) and aligned to the GRCh38 human reference genome by bwa v0.7.12 (Li and Durbin 2009). Mitochondrial reads and duplicate reads were removed and the remaining reads were filtered to include only properly aligned pairs by sambamba v0.7.1 (Tarasov, Vilella et al. 2015). Peaks were called from the deduplicated bam files using macs2 (Zhang, Liu et al. 2008) with parameters -q 0.001 --keep-dup all -B --SPMR --nomodel –nolambda. Peaks were merged across all four samples H3K9ac, H4K12ac, H3S10ph, or H4S1ph using bedtools v2.26.0 (Quinlan and Hall 2010) and intersected with refGene_hg38 gene models on the standard chromosomes. Modifications occurring in the gene body (excluding peaks found ±200 bp from transcription start sites) were counted and compared to classify genes with pan-acetylation only, pan-phosphorylation only, or both acetylation and phosphorylation. Genes with modifications occurring only within ±200 bp of their transcription start sites were similarly classified.

### Western Blotting (DRB inhibition and detection in HEK293 cell)

0.6 million HEK293T cells were seeded on the 6-well plate one day before treatment. On the next day, different inhibitors were added at the indicated concentration and incubated for 1 hour. Cells were then collected and lyzed with RIPA lysis buffer. The supernatant was collected by centrifugation and an equal amount of cell lysate was loaded on the SDS PAGE gel and transferred to the PVDF membrane for blotting analysis. Rat antibody 3E8 was used for the detection of serine5-phosphorated Pol II Rpb1 CTD. The membrane was then stripped with stripping buffer, and re-probed with mouse antibody 8WG16 for total Pol II Rpb1 CTD detection. Actin was used as a loading control.

### qPRO data process and figure generation

Figure 3b was generated per Hou et al., 2019 (Hou, Wang et al. 2019). In short, log2-comparative bigwig files were generated using DeepTools (ver. 3.5.4) bamCompare between control samples and each time point of treated samples. Reads with multiple alignments and PCR duplicates were ignored, and samples were scaled by RPKM. Comparative heatmaps were then generated from these log2-bigwig files using computeMatrix and plotHeatmap.

The annotation file was first split by strand, and filtered to only the maximally expressed isoform (highest average RPKM across all samples, minimum average of 0.1 RPKM) among all samples, and all genes whose TSSs were within a 100kb window another gene’s TSS on the same strand were identified and removed using the following command: `bedtools merge -i stdin -c 4,5,6 -o distinct,count,distinct | awk -F "\t" ’BEGIN{OFS="\t"} $5==1 {print $1,$2,$3,$4,$5,$6}’`. The two stranded annotation files were then recombined and sorted using `cat | bedtools sort -i stdin (bedtools ver. 2.28.0).` The annotation file was then used as input along with the comparative bigwig files as follows: computeMatrix reference-point --referencePoint TSS -b 5000 -a 100000 - R {annotation_file} -S {comparative bigwig files} --outFileName matrix.gz plotHeatmap -m matrix.gz -o heatmap.ref.svg --dpi 600 -- sortRegions ascend --sortUsing region_length.

### qPRO dataset generation- nuclei isolation

qPRO datasets were generated per Judd et al, 2020 (Judd, Wojenski et al. 2020). Samples were first treated and their nuclei isolated. In brief, cell cultures were collected in 50 ml Falcon tubes and centrifuged in a fixed-angle rotor centrifuge at 600 x g, 4 C for 5 minutes. The supernatant was removed, and ice-cold PBS (10 ml) was used to wash the pellet (wash repeated 3 times as above). The pellet was then resuspended in 10 ml ice-cold Lysis Buffer (10 mM Tris-HCl pH 7.5, 2 mM MgCl2, 3 mM CaCl2, 0.5% IGEPAL, 10% Glycerol, 2 UmL SUPERase-IN, brought to 10 ml with 0.1% DEPC-treated DI-water). The samples were then incubated for 10 minutes on ice. The samples were centrifuged in a fixed-angle rotor centrifuge at 1000 x g for 10 minutes at 4 C in 50 mL Falcon tubes. The resulting supernatant was poured off, and the pellet was resuspended with 1 ml lysis buffer, using a wide-mouth P1000 pipette tip. The volume of the sample was brought to 10 ml with lysis buffer, and centrifuged for 1000 x g, 4 C, 5 minutes (wash repeated as above twice). Nuclei were then resuspended in 1 ml freezing buffer (50 mM Tris-HCl pH 8.3, 5 mM MgCl2, 40% Glycerol, 0.1 mM EDTA pH 8.0, brought to volume with 0.1% DEPC treated DI-water), and transferred to a 1.7 ml Eppendorf tube. Nuclei were centrifuged at 1000 x g, 4 C, 5 minutes, and the supernatant was removed by pipetting. The pellet was resuspended with 500 uL freezing buffer. The nuclei were again pelleted at 1000 x g, 4 C, 5 minutes. The supernatant was removed, and the nuclei were resuspended in 110 uL freezing buffer, and stored at -80 C until library preparation.

### qPRO dataset generation- nuclei isolation

The qPRO protocol was adapted from Judd et al, 2020 (Judd, Wojenski et al. 2020). A detailed step-by-step protocol is available at dx.doi.org/10.17504/protocols.io.57dg9i6. In short, ice-cold isolated nuclei (100 ul) were added to 37 C 100 ul reaction buffer (Final Concentration: 5 mM Tris-Cl pH 8.0, 2.5 mM MgCl2, 0.5 mM DTT, 150 mM KCl, 10 units of SUPERase In, 0.5% sarkosyl, 20 uM rATP, 20 uM rGTP, 20 uM biotin-11-CTP, 20 uM biotin-11-UTP. The reaction was allowed to proceed for 10 min at 37 C. RNA was extracted twice with Trizol (500 ul), washed once with chloroform, and precipitated with 2.5 volumes of ice-cold ethanol and 1 uL GlycoBlue. The pellet was washed in 75% ethanol before resuspending in 30 uL of DEPC-treated water. The resulting RNA was denatured at 65 C for 30 seconds, and snap cooled on ice. RNA was then fragmented by base hydrolysis in 0.2 N NaOH on ice for 10–12 min, and neutralized by adding a 1× volume of 1 M Tris-HCl pH 6.8, and passed through a P-30 buffer exchange column (Bio-Rad #7326250). The 3’ adaptor was then ligated to the fragmented RNA (3’ RNA adaptor: /5Phos/GAUCGUCGGACUGUAGAACUCUGAAC/3InvdT/, final conc. 0.5 uM) with T4 RNA Ligase 1, and biotin-labeled products were enriched by a round of streptavidin bead enrichment (Dynabeads MyOne Streptavidin C1, Invitrogen #65001). For 5’ end repair, the RNA-bound beads were treated with T4 PNK (NEB #M0201S) and subsequently with RppH (NEB #M0356S). 5’ repaired RNA was ligated to reverse 5’ RNA adaptor (5’ UGGAAUUCUCGGGUGCCAAGG, final conc. 0.5 uM) before being extracted from the beads with trizol. RNA was then reverse transcribed into cDNA with SuperScript III (Thermo Fisher #12574026) and RT primer (5’ AATGATACGGCGACCACCGAGATCTACACGTTCAGAGTTCTACAGTCCGA). The product was amplified 15 +/- 3 cycles and products >150 bp (insert > 70 bp) using Phusion DNA polymerase (NEB #M0530S) and TruSeq Small RNA Library Preparation Kit barcoded primers. were size selected with 0.9X AMPure XP beads (Beckman #A63880) before being sequenced.

### Antibodies

Ser5ph-CTD of Pol II: mouse monoclonal antibody (3E8, Abcam cat.no.ab252852), 1:100; Native Rpb1 (native Pol II): mouse monoclonal antibody (8WG16, Santa Cruz cat.no.sc-56767), 1:100; Ser2ph-CTD of Pol II: mouse monoclonal antibody (3E10, Abcam cat.no.ab252855), 1:100; H3S10ph: mouse monoclonal antibody (CMA311, a gift from Dr. Hiroshi Kimura’s group), 1:100; H4S1ph: mouse monoclonal antibody (Abcam cat.no.ab177309), 1:100; H3K9ac: mouse monoclonal antibody (Active Motif. cat.no.61251), 1:100; H4K12ac: mouse monoclonal antibody (Active Motif.cat.no.39165), 1:100.

### Chemicals and Cell line

DRB, Medchem Express (cat.no.HY-14392); a-Amanitin, Medchemexpress (cat.no.HY-126683); CDK7 inhibitor LDC4297, Medchemexpress (cat.no.LDC4297); CDK8 inhibitor BI-1347, Medchemexpress (cat.no.BI-1347). Cell line: HEK293T, ATCC (cat.no.CRL-3216).

## Acknowledgments

We thank Drs. Philippa Marrack, Tony Gerber, Hua Huang, Marwan Abushawish, James Scott-Browne and other members of National Jewish Health for their suggestions and support. Drs. Carol C.L. Chen and Matthew C. Lorincz for suggestions and help. The work is partially supported by an NIH grant GM135421 (G.Z.) and funds from NB Life Laboratory LLC, specifically private support from Cheng-Yuan Zhang, Shi-Ning Xu, Lian-Hua Jin, Peng Sun, Yun-Xia Jiang, and Yongmei Jiang.

## Contributions

GZ conceived the concepts, designs of experiments, data analysis, and writing up the manuscript. LL and HL carried out most of the experiments. LL and SH for PRO-seq experiments. SH, SL, YZ, JG, QZ, and RD did the final data analysis and suggestions. TS and HK for H3S10ph antibody and suggestions.

## Supplementary Figures

